# Inhibitory control and the structural parcellation of the right inferior frontal gyrus

**DOI:** 10.1101/2020.08.13.249516

**Authors:** Rune Boen, Liisa Raud, Rene J. Huster

## Abstract

The right inferior frontal gyrus (rIFG) has most strongly, although not exclusively, been associated with response inhibition, not least based on covariations of behavioral performance measures and local grey matter characteristics. However, the white matter microstructure of the rIFG as well as its connectivity has been less in focus, especially when it comes to the consideration of potential subdivisions within this area. The present study reconstructed the structural connections of the three main subregions of the rIFG (i.e. pars opercularis, pars triangularis and pars orbitalis) using diffusion tensor imaging, and further assessed their associations with behavioral measures of inhibitory control. The results revealed a marked heterogeneity of the three subregions with respect to the pattern and extent of their connections, with the pars orbitalis showing the most widespread inter-regional connectivity, while the pars opercularis showed the least amount of connections. When relating behavioral performance measures of a stop signal task to brain structure, the data indicated a differential association of dorsal and ventral opercular connectivity with the go reaction time and the stopping accuracy, respectively.

## Introduction

The right inferior frontal gyrus (rIFG) is considered a key node for the inhibition of premature or no longer appropriate motor responses, which is one of the core aspects of behavioral flexibility and control (Aron et al., 2014, 2016; Swann et al., 2012). The IFG represents a structurally diverse area in the prefrontal cortex that usually is divided into three sub-regions based on its cytoarchitecture: the pars opercularis, pars triangularis, and pars orbitalis. Given that variability in the structural architecture of the brain often relates to specific aspects of behavior (Johansen-Berg, 2010), it is likely that the rIFG exhibits a richer functional diversity than often posited. A recent meta-analysis identified different functional clusters of the rIFG to be involved in distinct large-scale networks; only the posterior part (roughly corresponding to the pars opercularis) seemed to be involved in motor control, and was further divided into dorsal and ventral regions associated with response initiation and general inhibition, respectively (Hartwigsen et al., 2019). However, a structural connectivity map of rIFG subregions that would support this functional parcellation is lacking.

The rIFG has been suggested to be part of a right-lateralized fronto-basal ganglia network (Chambers et al., 2009; Jahanshahi et al., 2015), that instantiates inhibition of the motor cortex jointly with the pre-supplementary motor area (preSMA), the basal ganglia, and thalamic nuclei (Aron et al., 2014). Structural and functional connections have been established between the IFG, the preSMA (Swann et al., 2012), subthalamic nucleus (STN), and striatum (Isaacs et al., 2018). While the specific roles of the rIFG and the preSMA for response inhibition are not fully understood, increased fractional anisotropy (FA) in the pars opercularis has been negatively associated with inhibitory performance in a stopping task (i.e., shorter stop signal reaction times), while the reverse association has been reported for the preSMA (Xu et al., 2016). Thus, it seems possible that the rIFG and preSMA play complementary roles in the regulation of response inhibition. In order to fully understand the functionality of the stopping network, it is of fundamental importance to map the structural architecture of those regions that facilitate stopping of behavior.

The stop signal task (SST) is one of the most widely used paradigms to study response inhibition, and is often considered the most direct measure of reactive inhibition (van Belle et al., 2014), due to the possibility of calculating the stop signal reaction time (Logan et al., 1984). Yet, the SST additionally provides behavioral measures related to motor preparation under cognitive control, such as the trade-off between fast responding and accurate stopping, captured complementarily by the go reaction times (goRTs) and the stopping accuracy. This is important, because functional studies show that different rIFG subregions are involved in motor initiation as well as proactive and reactive inhibition (Hartwigsen et al., 2019; Messel et al., 2019). However, the interpretation of goRTs produced under the SST as task-general marker of motor preparation has been challenged. For instance, SST goRTs have been found to slow down with increasing probability of a stop signal (Zandbelt & Vink, 2010), which has been taken as evidence for a braking mechanism that proactively restrains responses (proactive inhibitory control) (Albares et al., 2014; Zandbelt & Vink, 2010). Thus, motor initiation in the SST seems to be influenced by other cognitive mechanisms, such as strategic slowing in order to balance performance speed and accuracy (Leotti & Wager, 2010). Correspondingly, it has been found that activations associated with go responses in the SST overlap with those related to outright stopping (e.g., the preSMA and striatum, Forstmann et al., 2008), while others have reported that the interplay between the IFG and preSMA is involved in response slowing (Gaal et al., 2010).

Contrasting the SST with a response choice task represents the ideal tool to study the associations of rIFG subdivisions with respect to their potential involvement in response generation and inhibition. We therefore investigated the associations of rIFG subregions with response initiation (responding without stopping constraints in a pure response choice task), response initiation under proactive inhibitory control (goRT in SST), as well as response inhibition under reactive inhibitory control (stop signal reaction time and accuracy in the SST). The primary aim of the present study was to map the structural connections of three subregions of the rIFG: the pars opercularis, pars triangularis and pars orbitalis. Further, we extended the abovementioned literature by investigating the white matter fiber pathways from the dorsal and ventral region of the pars opercularis to regions critical for motor control. We expected that the dorsal and ventral connections would show differential functional associations such that connections from the dorsal part would be associated with response initiation, while those of the ventral part would show associations with measures of response inhibition.

## Methods

### Participants

Thirty-one participants took part in the experiment (14 females, mean age = 26.35, range = 20-36 years). One participant was excluded from behavioral and connectivity analyses due to technical issues that caused partial data loss. Five participants were excluded from the behavioral analyses: two participants were excluded due to technical issues with the response device, two more did not complete the behavioral tasks, and one participant was excluded due to an interruption in the middle of the experiment, leading to non-convergence of the stop signal delays (SSD). This resulted in 30 participants for the structural connectivity analyses and 25 participants for the analysis of brain-behavior associations. All participants were right-handed, had normal or corrected to normal vision and reported no history of psychiatric or neurological disorders, migraine, or loss of consciousness. The experiment was approved by the internal review board of the Department of Psychology, University of Oslo. All participants gave informed consent and received a gift card of 300 NOK for participation.

### Image acquisition

All magnetic resonance imaging (MRI) sequences were run on a 3.0 Tesla Philips Ingenia whole-body scanner (Philips Medical Systems, Best, the Netherlands) with a 32-channel head coil. Diffusion-weighted imaging (DWI) was performed using a single-shot EPI sequence, one b0 image, and diffusion weighting was conducted across 32 non-collinear directions with a b-value = 1000s/mm2, flip angle = 90 degrees, repetition time (TR) = 13.45s, echo time (TE) = 62ms, field of view (FOV) = 224 x 224 x 120, Matrix = 96 x 94 x 60. The acquired voxels of size 2.33 x 2.38 x 2.0 mm were reconstructed to 2.0 mm isotropic voxels. T1 images were acquired using the following parameters: TE =2.3, TR = 5.1, FOV = 256 x 256 x 184, Matrix = 256 x 254 x 184, voxel size = 1.0 x 1.0 x 1.0 mm.

### Data processing

All processing steps were conducted in ExploreDTI v.4.8.6 (Leemans, Jeurissen, Sijbers, & Jones, 2009). All images were inspected for artifacts and excessive head movements, corrected for eddy current-induced distortions and head motions with a non-diffusion weighted image as reference. Each participant’s high-resolution T1-weighted image was utilized for EPI correction, and the DWI data were correspondingly resampled and transformed into 1mm isotropic voxels.

### Brain atlas and tractography

A standardized brain atlas consisting of the Automated Anatomical Labeling (AAL) atlas (Tzourio-Mazoyer et al., 2002) and a bilateral binarized mask of the STN (Forstmann et al., 2012) were used to outline 92 brain regions across both hemispheres. The preSMA and SMA were not separated due to tracts ending in the border of preSMA and SMA (Catani et al., 2012), and will collectively make up a region to be referred to as SMA complex (SMAc). A whole brain deterministic tractography with every voxel as seed point was completed and utilized to map the connectivity of ending and passing tracts of the three sub-regions of the rIFG. An ending connection was determined between two regions if the reconstructed fiber pathway originated in one of the regions and terminated in the other. A connection was deemed a passing pathway if the reconstructed tract passed through the regions. Seed point resolution was set to 1mm isotropic, with an FA threshold of 0.2 and an angle threshold of 45 degrees. We reran the same procedure after segmenting the pars opercularis into a dorsal and ventral region based on a halfway split along its longest extent.

### Tasks and procedure

All participants were measured on two separate days (with a median interval of 1 day). Session one consisted of three MRI sequences, including a T1, DWI and resting-state fMRI measurement. Session two consisted of a concurrent measurement of electroencephalography (EEG), single-pulse transcranial magnetic stimulation (TMS) and electromyography (EMG) during two separate computer-based experiments: the delayed response task (DRT) and the stop signal task (SST). As this study focused on the associations of white-matter structure with behavior, the acquired EEG, EMG, and TMS data will not further be regarded here.

The experimental tasks were developed as in-house MATLAB scripts (The MathWorks, Inc., Massachusetts, USA) using the Psychophysics Toolbox (Brainard, 1997; Kleiner et al., 2007; Pelli, 1997). Participants sat in a chair at a viewing distance of 1 meter from the monitor and responded on separate response devices with their left and right index fingers. The screen resolution was 1280*1024 with a refresh rate of 60Hz. The experimental tasks consisted of a cued DRT of 3 blocks and a cued SST of 12 blocks. Each block took approximately 6 minutes to complete with the possibility to take breaks of self-determined durations in-between each block and task (total time = 92,4 minutes + pauses). Trials containing TMS pulses were excluded from the analyses. The DRT data consisted of 96 non-TMS pulse trials with 72 go-trials and 24 catch trials, while the SST consisted of 432 non-TMS pulse trials with 288 go-trials and 144 stop trials. The go and stop stimuli were presented as circles colored either blue or orange. The colors of the stimuli were counterbalanced for the go and stop signal; the color of the go signal remained the same for the DRT and SST throughout the experiment.

The cued DRT started with a fixation cross randomly jittered between 1800-2300 ms. After this, a cue (i.e. a right or left leaning bracket) was presented that indicated which finger to prepare for a response (e.g., right leaning bracket = right index finger). The inclusion of these valid cues eliminated the decision making phase after the detection of the go signal (as the decision about which hand to use is shifted to the cue-delay period), and thus allows for the investigation of response initiation without confounding response conflict. The cue duration was fixed at 900 ms. The go signal (a circle next to the bracket) appeared after the cue and was present for 800 ms or until a response was made. A go signal was omitted in 9% of the trials to diminish premature responding. The SST was similar in all aspects of the task but two: i) the inclusion of a stop signal in a minority of the trials, and ii) that no go signals were omitted. Stop signals appeared in 33% of the trials and were presented after a stop signal delay (SSD) that was adjusted following a tracking procedure. The SSD was initially set to 250 ms for both hands and was subsequently adjusted based on the performance in the preceding trial. The SSD was increased by 33 ms if the previous stop signal trial was successful and decreased by 33 ms after unsuccessful stop trials. The minimum and maximum SSD were set to 80ms and 800ms, respectively.

### Instructions

For the DRT, participants were told to respond as fast as possible to the circle appearing next to the cue. For the SST, the participants were told that the task was similar to the DRT, but that a stop signal would be shown on a minority of the trials to which they should try to withhold their response. They were further instructed to be as fast and accurate as possible and that mistakes were to be expected during the task. In go trials, feedback (“too late”) was presented if no response was produced within 800 ms after the go signal. The participants were also shown feedback after each block. If the average goRT of the preceding block was above 600ms, the participants were instructed to be faster. However, if the average accuracy was below 45%, they were instructed to be more accurate. If the participants’ performance was within these thresholds, they were presented with the feedback “Well done”.

### Derivation of dependent variables and statistical analyses

To quantify white matter microstructure, we extracted the FA values of the tracts of interest from the whole brain tractography analyses. Further, the average FA across the brain for each participant was derived by calculating the mean value of the FA for all passing and ending tracts across the brain and averaging these into a single global FA value. To test if the rIFG subregions differed in their connections to other brain regions, we calculated the total number of reconstructed connections from each subregion for each participant. Subsequently, we ran paired t-tests with the number of reconstructed connections detected from these rIFG subregions.

The following behavioral measures were extracted from the DRT and SST: Go-accuracy, goRT, probability of choice errors, omissions, and premature responses (responses given after the cue, but before go signal onset). For the SST, we also calculated the stopping accuracy, unsuccessful stop RT, stop signal delay, and stop signal reaction time (SSRT). The SSRTs were estimated based on the integration method (Verbruggen & Logan, 2009). Specifically, the goRT distribution for each participant was extracted that included premature responses and go errors, and the omissions were replaced by the maximum go RT (Verbruggen et al., 2019). The SSRT was calculated by subtracting the mean SSD from the nth value in the sorted goRT distribution, where n corresponds to the probability of responding in the stop trials multiplied with the number of values in the go RT distribution. All behavioral measures are reported as an average of both hands. The association between the goRT and SSRT in the SST was calculated as a parametric bivariate correlation. All statistical analyses assessing behavioral and brain-behavior associations were carried out with IBM SPSS Statistics for Windows, Version 25.0.

### Brain-behavior analyses

The goRT, SSRT and stopping accuracy were used as dependent variables, and the global FA and parameters of the tracts from the dorsal and ventral part of the pars opercularis and to the target region SMAc were used as predictor variables in the regression analyses. The global FA was included to account for overall inter-individual differences in white matter microstructure of the brain. We specifically focused on the pars opercularis as it has been considered the key node of inhibitory control (Aron et al., 2014; Hartwigsen et al., 2019). For visualizations and brain-behavior analyses, we used tracts that were present in at least 80% of the participants for generalizability and reliability.

## Results

### Structural connectivity maps of the three rIFG subregions

The structural connections of the rIFG sub-regions are visualized in Figure 1 and the average number of passing and ending tracts for each subregion is depicted in Figure 2. In total, the three subregions showed extensive connections that covered all four lobes in the right hemisphere, as well as several structures within the basal ganglia. The pars opercularis (Fig.1A) exhibited a similar connectivity pattern as the pars triangularis, albeit with fewer connections (Fig.1B), while the connectivity fingerprint of the pars orbitalis (Fig.1C) exhibited a more widespread network that also reached peripheral regions such as the occipital cortex. The data therefore suggest a posterior to anterior gradient with increasing connectivity from the opercularis, via the triangularis, to the orbitalis. To quantitatively test this observation, we computed pair-wise t-tests between these regions with the number of connections estimated for each subject as dependent variable (Figure 2). These tests were run separately for both passing and terminating projections. The results revealed a significantly lower amount of terminating connections for the pars opercularis compared to the pars triangularis (t (29) = −6.67, p < .001) and the pars orbitalis (t (29) = −9.04, p < .001), while the pars triangularis showed fewer projections compared to the pars orbitalis (t (29) = −3.98, p < .001). A similar pattern emerged for passing connections, where the pars opercularis had fewer passing projections compared to the pars triangularis (t (29) = −8.46, p < .001) and pars orbitalis (t (29) = −14.38, p <.001), while the pars triangularis exhibited fewer connections compared to the pars orbitalis (t (29) = −6.59, p < .001).

**Figure 1.**
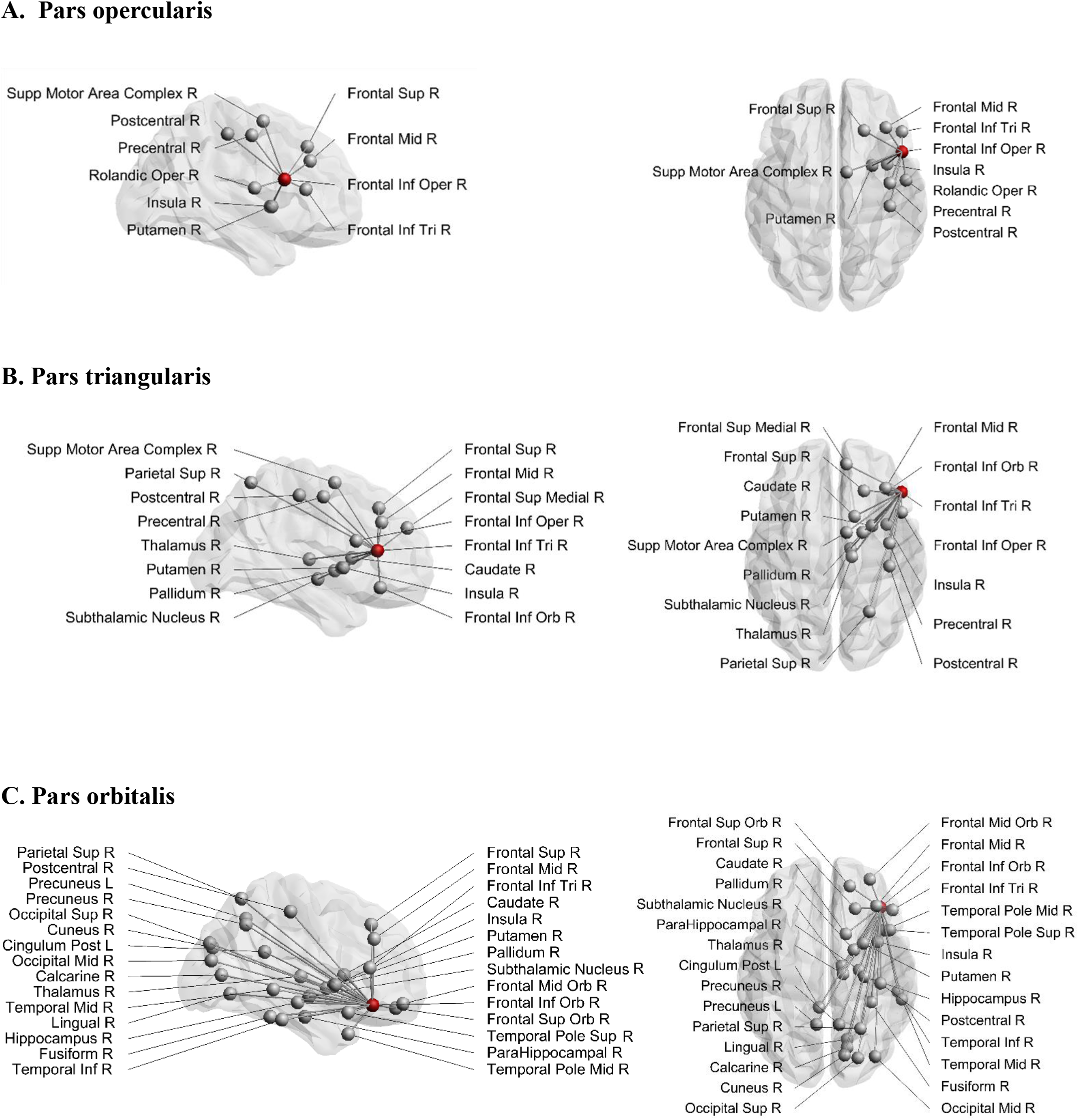
Structural connections from A, pars opercularis, B, pars triangularis, and C, pars orbitalis. The seeding region is marked as a red node. Sup = superior, Inf = inferior, Mid = middle. Supp = supplementary, Oper = opercularis, Tri = triangularis, Orb = orbitalis, R = right, L = left.

**Figure 2.**
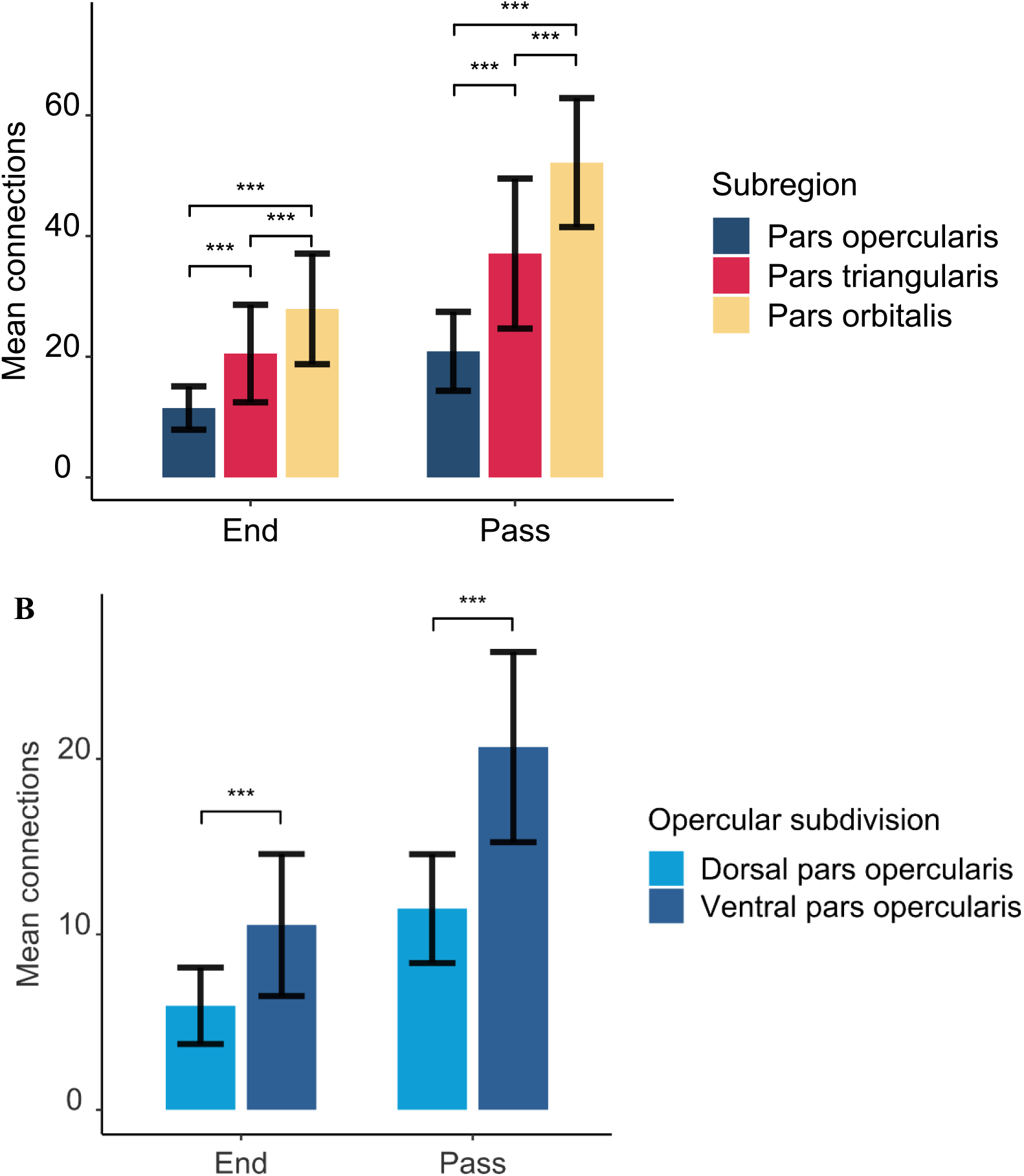
The average number of connections seeding from A) the rIFG subregions and B) opercular subdivisions. End and pass represent whether the tracts were part of an ending or passing fiber pathway. The error bars represent standard deviations. The asterisks above the bars mark the significance level at p < .001.

### Structural connectivity maps of the dorsal and ventral pars opercularis

The structural connections of the dorsal and ventral pars opercularis are visually presented in Figure 3. While the connectivity patterns of these two subregions show considerable overlap, the ventral part of the pars opercularis exhibited more connections as indicated via significant paired t-tests for both ending (t(29) = −6.49, p < .001) and passing (t(29) = −9.20, p < .001) tracts. Connectivity differences emerged such that the dorsal opercularis showed a connection to mid-frontal cortex, whereas the ventral opercularis showed connections to postcentral cortex, rolandic operculum, insula and putamen.

**Figure 3.**
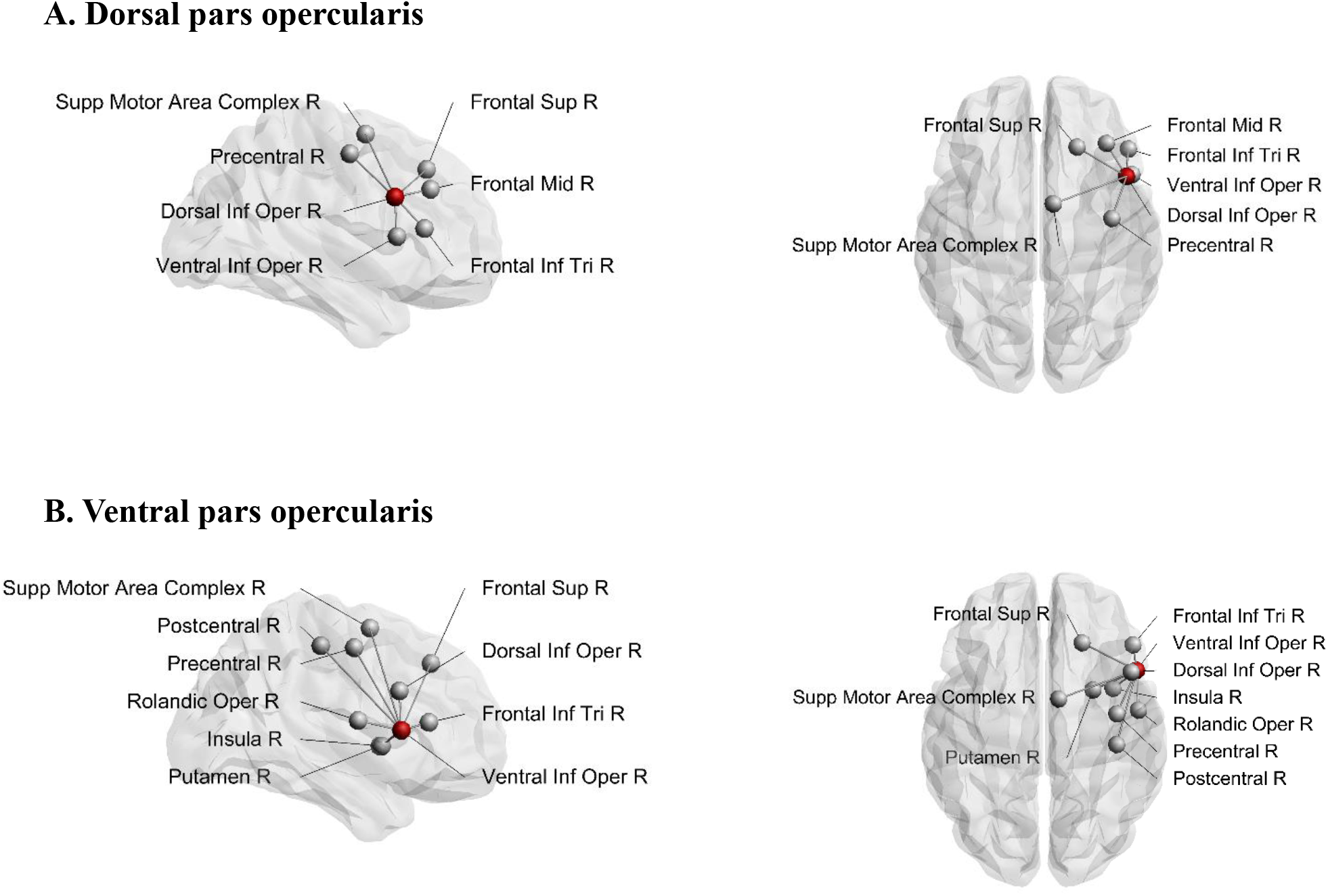
Structural connections from A, dorsal pars opercularis, and B, ventral pars opercularis. Seeding region is marked as a red node. Sup = superior, Inf = inferior, Mid = middle, Supp = supplementary, Oper = opercularis, Tri = triangularis, R = right

### IFG connectivity within the stopping network

We conducted a connectivity analysis that specifically focused on differential connectivity patterns of the three IFG subregions with the other brain areas considered part of the stopping network: the SMAc, insula, caudate, putamen, and the STN. Connections to the stopping network were deemed to be reliably present if they were identified in at least 80% of the participants. Figure 4 depicts the average frequencies of these connections and Figure 5 depicts the connections. The pars opercularis showed reliable connections to the SMAc, insula, and putamen. The pars triangularis showed reliable connections to the SMAc, insula, putamen, caudate, and the STN. The pars orbitalis exhibited reliable connections to the insula, putamen, caudate and the STN. Thus, the three rIFG subregions showed a differential connectivity within the stopping network, with connectivity in pars opercularis being limited to the cortical areas and putamen, while the other two regions showed additional subcortical basal ganglia connections.

**Figure 4.**
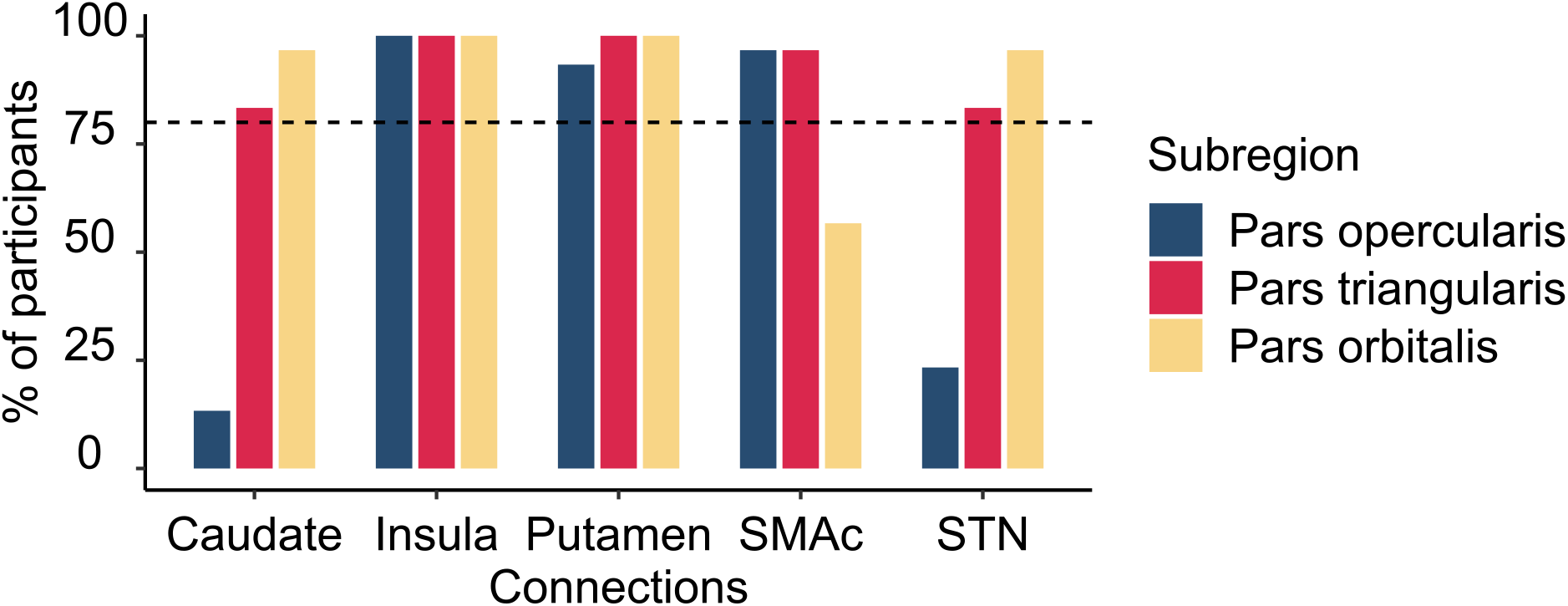
Histogram illustrating the percentage of participants having connections from the rIFG subregions to the regions within the stopping network. The vertical dashed line refers to the inclusion threshold of 80% for a reliable connection. SMAc = supplementary motor area complex, STN = subthalamic nucleus.

**Figure 5.**
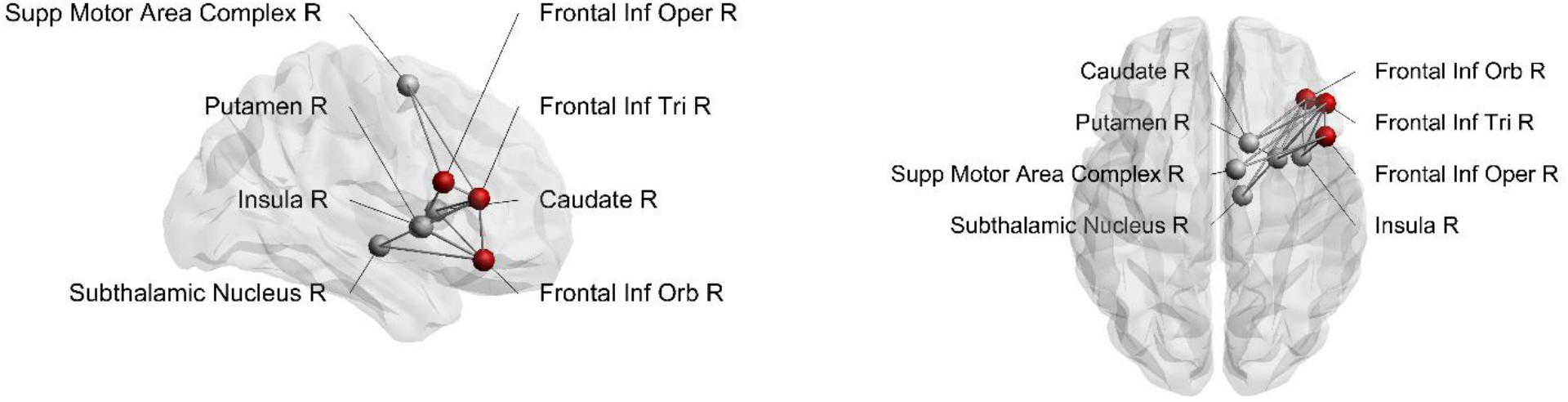
Structural connections from rIFG subregions to the stopping network. Seeding region is marked as a red node. Inf = inferior, Supp = supplementary, Oper = opercularis, Tri = triangularis, Orb = orbitalis, R = right

### Behavioural results

Descriptive statistics of the behavioural measures obtained from the DRT and SST are presented in Table 1. Across participants, the average accuracy (≥ 95% in both tasks) indicated good task performance. The average stop accuracy was 48%, which indicated successful SSD tracking, and all participants showed faster unsuccessful stop RTs than go RTs. The goRTs were shorter in the DRT than in the SST (t (24) = −9.18 p < .001) and did not correlate with each other (r = .081, p = .700). The average SSRT was 209 ms and did not correlate significantly with the mean goRT in the SST (r = −.30, p = .15). However, stopping accuracy showed a significant association with goRT (r = .57, p = .003) and SSRT (r = −.60, p < .001) in the SST.

**Table 1.** Behavioral characteristics

|  | DRT | SST |
| --- | --- | --- |
| Go accuracy, % | 96 (3.92) | 95 (2.7) |
| Choice errors, % | .38 (.74) | .24 (.34) |
| Go omissions, % | 1.48 (1.73) | 3.15 (1.62) |
| Premature responses, % | 1.89 (3.35) | 1.21 (1.68) |
| Go RT, ms | 320 (53) | 475 (70) |
| Stop accuracy, % | - | 48 (3.83) |
| Unsuccessful stop RT, ms | - | 409 (74) |
| Stop signal delay, ms | - | 293 (91) |
| Stop signal reaction time, ms | - | 209 (26) |

*Note.* Go RT = mean reaction time on go trials, ms = milliseconds, standard deviations are presented in the brackets.

### Global and tract-specific associations with behavior

First, we tested whether the global FA was predictive of task performance, and found that the global FA value was not significantly correlated with the DRT goRT (r =.091, p = .664), but that it exhibited significant correlations with the SST goRT (r = .434, p = .030), SSRT (r = −.414, p = .040) and stopping accuracy (r = .479, p =.015).

We then focused more specifically on key regions of the stopping network. Given the putative interactions of the pars opercularis and the SMAc in the stopping literature and their role in motor and inhibitory control, we computed a linear regression analysis using the FA of the dorsal pars opercularis-SMAc and the ventral pars opercularis-SMAc tracts as predictors of DRT goRT, SST goRT, SSRT, and stopping accuracy (Table 2). The global FA was added as a covariate, given the aforementioned associations of global FA with task performance measure, to further test the regional specificity of the described effects. The full model was only significant for the SST goRT and accuracy with the dorsal pars opercularis-SMAc tract as a significant predictor. The partial regression plots of the FA of the dorsal pars opercularis-SMAc tract predicting goRT and stopping accuracy in the SST are presented in Figure 6.

**Table 2.**
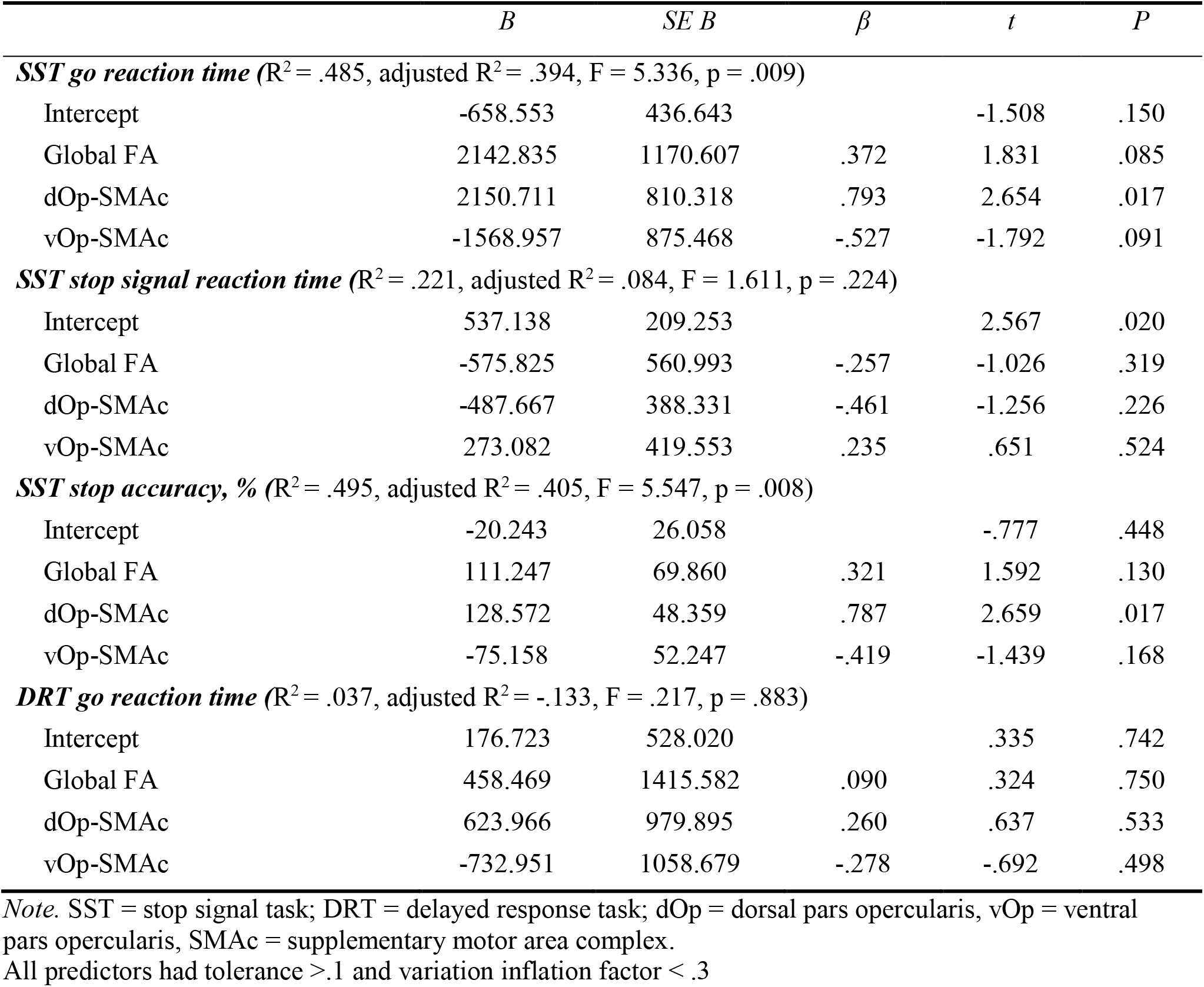
Summary of Multiple Regression Analyses (N =20)

**Figure 6.**
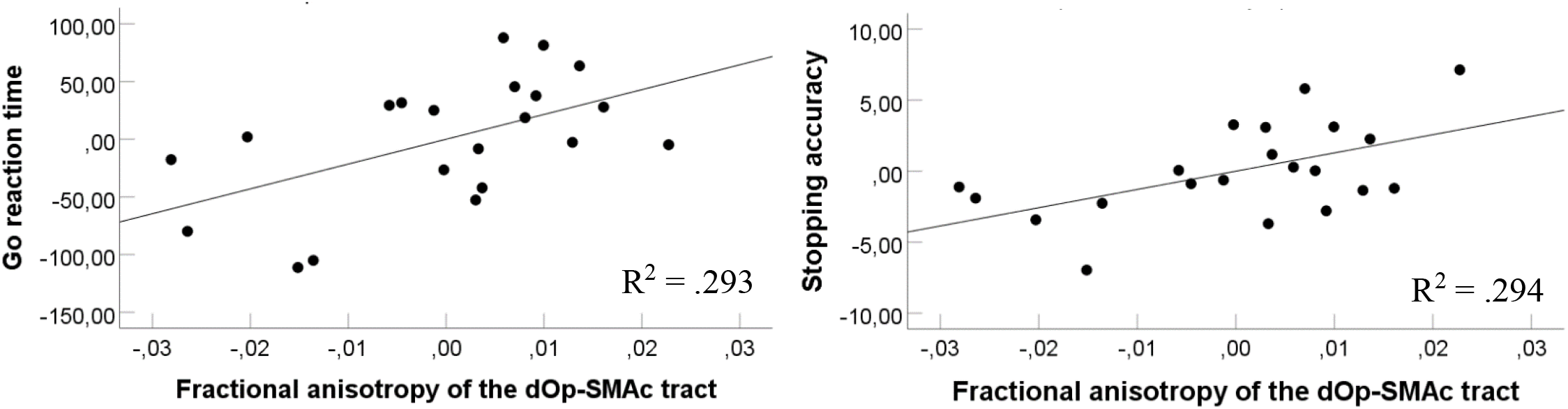
Partial regression plots of the fractional anisotropy of the dOp-SMAc tract predicting go reaction time (left) and stopping accuracy (right) in the stop signal task, controlling for whole brain fractional anisotropy and the ventral pars opercularis-SMAc. Both correlations were significant at p < .05. dOp = dorsal pars opercularis, SMAc = supplementary motor area complex.

## Discussion

Our primary objective was to investigate the white matter fiber pathways of three rIFG sub-regions (i.e. pars opercularis, pars triangularis and pars orbitalis) using diffusion weighted imaging and deterministic tractography. The three subregions showed substantial differences in their connectivity patterns, as well as a posterior to anterior gradient in the amount of connections. In addition, the pars opercularis was segmented into a dorsal and ventral region, both of which were shown to have connections to SMAc. However, only the fractional anisotropy of the dOp-SMAc tract was a significant predictor of task behavior, namely for the goRT and stopping accuracy in the SST.

Hartwigsen and colleagues (2019) identified functionally diverse subregions in the rIFG, following a posterior-to-anterior axis, where the posterior part was associated with motor functioning and the anterior part was related to abstract cognitive functions. In relation to this, we found evidence for a posterior-to-anterior division of structural connections within the rIFG. That is, the pars orbitalis showed the highest amount of connections, followed by the pars triangularis, while the pars opercularis exhibited the lowest amount of connections. Moreover, the connectivity fingerprints of the pars opercularis and pars triangularis were largely restricted to central and frontal regions, while the pars orbitalis showed the most widespread inter-regional connectivity pattern among the three rIFG sub-regions. This is interesting as the pars orbitalis has been associated with abstract cognitive functions (Hartwigsen et al., 2019), whereas the current study indicates that it also shows a widespread connectivity pattern reaching regions across all four lobes in the right hemisphere. Speculatively, it might be that the widespread connections of the pars orbitalis serve its involvement in complex cognitive functioning, such as abstract thinking and social cognition. This is in contrast to the posterior part of the rIFG, which has been proposed to be crucial for inhibitory control (Aron et al., 2014), with a further subdivision of a dorsal region involved in motor execution and a ventral region involved in motor inhibition (Hartwigsen et al., 2019). In the current study, the segmentation of the pars opercularis into a dorsal and ventral region revealed some marked differences where the ventral part of the pars opercularis showed a higher inter-regional connectivity compared to the dorsal part. In addition, both regions exhibited connections to the SMAc, an important region within the stopping network. Altogether, in line with previous evidence suggesting a functional divergence in the rIFG along its posterior-to-anterior axis, we found further evidence that the rIFG subregions are also structurally diverse along the posterior-to-anterior axis.

We also identified several connections of the rIFG sub-regions to other parts of the stopping network. The pars opercularis showed reliable connections to the SMAc, insula, and putamen.

Surprisingly, we did not find evidence for a reliable connection from the pars opercularis to the STN. This is in contrast to previous research that has shown this connection (Isaacs et al., 2018), albeit with data acquired with ultra-high field MRI and probabilistic tractography. However, the current results do show a reliable connection from both the pars triangularis and pars orbitalis to the STN. This might indicate that a connection between the pars opercularis and the STN consists of a tract with a complex architecture, which is harder to reconstruct with the conservative tractography technique used in the present study. It is interesting to note, however, that the pars triangularis was the only rIFG subregion that showed a reliable connection to the SMAc, insula, putamen, caudate and STN. Given the overlapping connectivity fingerprints of the pars opercularis and pars triangularis, the combination of these regions might be a more suitable connectivity hub for inhibitory control compared to pars opercularis alone.

Furthermore, Hartwigsen and colleagues (2019) suggested that the posterior part of the rIFG could be segmented into a dorsal and ventral region and that these regions are associated with motor initiation and inhibition, respectively. However, it is unclear whether the dorsal part relates to the cognitive effort necessary to execute correct responses in demanding tasks, or whether it relates to motor execution proper. In the current study, both the dorsal and ventral regions of the pars opercuarlis showed connections to the SMAc, a connection that has been suggested to be important for inhibitory control (Aron et al., 2007; Swann et al., 2012). Thus, it is interesting that our results revealed a significant positive relationship between the dorsal pars opercularis-SMAc and the goRT from the SST, while the ventral pars opercularis-SMAc showed a (non-significant but considerable) negative relationship with the goRT. We also observed the same pattern for the stopping accuracy, showing that increased connectivity strength in the dorsal pars opercularis-

SMAc is related to increased reaction time and stopping accuracy. This is interesting in context of previous research that showed increased fractional anisotropy in the pars opercularis to be negatively associated with the SSRT, while increased fractional anisotropy in the preSMA was positively associated with the SSRT(Xu et al., 2016). Moreover, the dorsal pars opercularis-SMAc tract was a significant predictor of goRT in the SST and not the DRT, and the goRTs from the DRT and SST did not correlate. This suggests that the goRTs obtained from the SST are influenced by other cognitive control mechanisms than motor generation alone. This supports a role of the dorsal opercularis in cognitively demanding motor initiation or the balancing of response speed and accuracy as opposed to plain motor generation in itself. The observed pattern thus supports the hypothesis of different functional roles of the dorsal and ventral parts of the opercularis.

In conclusion, the results indicate that the three sub-regions of the rIFG exhibit heterogeneity in terms of their connectivity, which is supported by the difference in the intra and inter-individual amount of tracts across the sub-regions. The overall pattern followed a posterior to anterior gradient with increasing connectivity from the pars opercularis, via the pars triangularis and to the pars orbitalis. Although, the pars orbitalis showed the most widespread connectivity, all three rIFG subregions showed several connections to regions implicated in inhibitory control. The segmentation of the dorsal and ventral pars opercularis showed that both regions had reliable connections to the SMAc, but only the ventral part was connected to the insula and putamen. Finally, the brain-behavior associations further supported a functional differentiation between the dorsal and ventral pars opercularis, implicating them in response execution under increased cognitive control and inhibition, respectively.

## Notes

Conflicts of interest: None

### Competing Interest Statement

The authors have declared no competing interest.

